# APP-KI mice do not display the hallmark age-dependent cognitive decline of amyloid diseases

**DOI:** 10.1101/2024.05.24.595745

**Authors:** Lisa Blackmer-Raynolds, Lyndsey D. Lipson, Isabel Fraccaroli, Ian N. Krout, Jianjun Chang, Timothy Robert Sampson

**Author notes:** These authors contributed equally (Lisa Blackmer-Raynolds, Lyndsey D. Lipson).

## Abstract

APP knock-in (KI) mice serve as an exciting new model system to understand amyloid beta (Aβ) pathology, overcoming many of the limitations of previous overexpression-based model systems. The APP^SAA^ mouse model (containing the humanized APP with three familial Alzheimer’s disease mutations) and the APP^WT^ control are the first commercially available APP KI mice within the United States. While APP^SAA^ mice have been shown to develop progressive Aβ pathology and neuroinflammation, behavioral changes, particularly in cognitive functions, have yet to be described. Therefore, we performed an in-depth longitudinal study over 12 months, assessing cognition in these two strains, as well as assessments of motor and GI function. We surprisingly note no overt, progressive cognitive impairment or motor deficits. However, we do observe a significant increase in fecal output in APP^SAA^ mice compared to APP^WT^ at 12 months old. These data provide a baseline for these models’ behavioral attributes.

**Highlights:**

- APP^SAA^ and APP^WT^ knock-in mice do not display age related cognitive decline
- Fecal output appears altered by APP genotype, but no other measure of GI function is impacted.
- Both genotypes behave equally in motor function tests

## Introduction

Establishing and characterizing animal models that accurately recapitulate pathological and behavioral outcomes of human disease remains an essential component—and limiting factor—to understanding disease etiology and identifying novel treatment targets. Rodent models of amyloid beta (Aβ) pathology—a key hallmark of Alzheimer’s disease (AD), as well as a prominent co-pathology in Parkinson’s disease (PD), Lewy body dementia (LBD), and other amyloid diseases—have been utilized for decades ^1,2^. These models have provided important insights into the role of Aβ in neurodegeneration and other physiological processes. Early rodent models of Aβ pathology were first developed by overexpressing the human amyloid precursor protein (APP) gene containing highly amyloidogenic mutations associated with familial AD (FAD) in humans ^3-5^. The majority of amyloid transgenic mouse models in use today follow a similar, overt overexpression approach, driving high expression of various combinations of transgenes of APP, presenilin, tau, and alpha-synuclein ^1,6^. However, there is increasing recognition that these overexpression approaches also have a number of important limitations. These include random gene integration, ectopic expression, overrepresentation of specific splice variants, non-physiological drivers of protein expression, and an inability to assess transcriptional regulatory impacts on the gene of interest, all of which confound experimental results ^7^.

In response to these limitations, various knock-in (KI) models of amyloid pathology have been created in which the murine gene of interest is knocked-out and replaced with a humanized transgene either at the locus itself or in *trans* ^1,8-15^. These KI models overcome many of the limitations of overexpression based models because the transgenes are expressed closer to physiological levels and are driven by native promotors, generating much excitement within the research community ^7^. The first commercially available KI model of Aβ pathology in the United States, the APP^SAA^ mouse, carries humanized APP with three mutations associated with familial AD (the Swedish, Arctic, and Austrian mutations) ^15^. The APP^SAA^ mouse develops amyloid pathology, neurodegeneration, and neuroinflammation that recapitulates elements of human disease ^15^. However, it is currently unknown whether these pathological hallmarks are associated with cognitive behavioral outcomes consistent with disease progression in humans. To address this, here we performed an in-depth, longitudinal characterization of various aspects of rodent behavior known to be impacted by amyloid pathology including anxiety-like behaviors, learning and memory, as well as motor and gastrointestinal function in both the APP^SAA^ and APP^WT^ mouse (with the wildtype human APP gene knocked-in) lines. We predicted that both strains would display progressive behavioral impairments, and that APP^SAA^ mice would comparatively show exacerbated impairments. Despite significant human Aβ accumulation in the brain of APP^SAA^ mice, neither strain was found to develop progressive cognitive or motor deficits, even at 12 months of age. However, APP^SAA^ mice were found to display elevated fecal output compared to APP^WT^ mice, at 12 months old, suggesting potential impacts of the SAA genotype on GI functions.

## Materials and methods

### Animal husbandry

Female and male APP^SAA^ KI (B6.Cg-Apptm1.1Dnli/J Strain #:034711) and APP^WT^ KI (B6.Cg-Appem1Adiuj/J Strain #:033013) mice were acquired from Jackson Labs through Dr. Srikant Rangaraju and maintained as homozygotes. Mice were genotyped by PCR using primers and conditions per the vendor (Jackson Labs). Mice were housed within a central, specific pathogen-free vivarium in microisolator cages on ventilated racks, with food (LabDiet: 5001) and water provided *ad libitum* and a 12:12hr light-dark cycle. All animal experiments were approved by the Institutional Animal Care and Use Committee of Emory University (PROTO201900056).

### Cognitive behavior assessments

Cognitive testing was performed longitudinally when mice were 2, 4, 5, 6, 8, and 12 months old. *Open Field* (OFT): Mice were placed in a 45cm square open field box for 10 minutes while distance traveled and time spent in center was recorded. *Object location test* (OLT): 24 hours post OFT, the OLT was run as described in dx.doi.org/10.17504/protocols.io.rm7vzxdo4gx1/v1. Briefly, testing consisted of a 10-minute study phase, 10-minute retention delay, and 5-minute testing phase. Object exploration was considered time with nose within 2cm of an object. Exploration ratio = moved object exploration/total object exploration. *Y-maze:* Mice were then tested on the Y-maze for 8 minutes, as described in dx.doi.org/10.17504/protocols.io.eq2lyjr1mlx9/v1. Percent alternation = total alternations (consecutive entries into 3 arms) / maximum possible alternations (total entries).

### Barnes Maze

Barnes maze testing was adapted from^16^, and performed as described in dx.doi.org/10.17504/protocols.io.kxygx3bozg8j/v1. Testing occurred on a 92cm diameter, 20 hole, Barnes maze (MazeEngineers) over a 6-day period, with one habituation trial, five 3-minute training trials, and a 72-hour probe trial. Primary latency and time spent in target quadrant were recorded for each training and probe trial. All behavioral tracking and analysis was performed using EthoVision XT software (Noldus Information Technology, Wageningen, the Netherlands) and the testing arenas/objects were cleaned between trials with 70% ethanol to eliminate olfactory cues.

### Motor behavior assessments

Motor function testing occurred at the experimental endpoint of 12mo of age as follows. *Adhesive removal:* The adhesive removal test was performed as described in dx.doi.org/10.17504/protocols.io.q26g7pjo9gwz/v1, by placing an adhesive sticker onto the mouse’s nose and quantifying the time needed to remove the sticker across three consecutive trials. *Wirehang:* The wirehang test was performed as described in dx.doi.org/10.17504/protocols.io.3byl4qy9zvo5/v1. Briefly, mice were placed on a 43cm square wire screen (made up of 12 mm squares of 1 mm diameter wire), turned upside down, and the time the mouse held onto the screen was recorded and averaged across three consecutive trials. *Hindlimb rigidity:* Hindlimb scoring was performed on a scale of 0-3 based on extent of hindlimb clasping as described in doi.org/10.17504/protocols.io.n2bvj3mnnlk5/v1 on two separate days by two independent scorers ^17^.

### Intestinal behaviors

At the experimental endpoint of 12mo of age, mice were assessed for general GI function through the following behaviors. *Fecal output:* Fecal output was performed as described in dx.doi.org/10.17504/protocols.io.rm7vzj3j5lx1/v1 by placing mice in a 1L plastic beaker and recording the number of fecal pellets cumulatively every 5 minutes for 30 minutes. *Total intestinal transit:* Intestinal transit time was measured using carmine red dye elution as described in dx.doi.org/10.17504/protocols.io.eq2lywpwwvx9/v1. Mice were gavaged with 100μl of sterile carmine red dye (6% w/v) (Sigma, C1022) dissolved in 0.5% methylcellulose (Sigma, M7027) and checked for elution of a red pellet every 15 minutes for 8 hours.

### Tissue Collection and Aβ Quantification

At 12 months of age, mice were humanely euthanized under isoflurane anesthesia and perfused with PBS. Hippocampal tissue was dissected, flash frozen, and protein extracted using a three-part fractionation protocol dx.doi.org/10.17504/protocols.io.j8nlk8ky1l5r/v1. Tris soluble (1M Tris HCL, 0.5M MgCl2 and 0.1M EDTA-pH 7.8) and Triton soluble (1% Triton-X100) fractions were run on the Meso Scale Discovery V-PLEX human Aβ peptide kit (K15200E) according to the manufacture’s guidelines and the resulting protein values were normalized to frozen tissue weight.

### Statistical Analysis

As indicated in the figure legends, data are expressed as mean ± s.e.m. Statistical tests were performed using GraphPad Prism 8. All raw numerical data are included with the manuscript as a supplemental file.

## Results

In order to test the hallmark, progressive, cognitive impairments observed in amyloid diseases, and other amyloid mouse models, we examined both APP genotypes across a battery of behavioral tests. Firstly, in the open field test (OFT) we observe a gradual decrease in distance traveled over time-a common finding as animals become habituated to the repetitive testing environment-which is qualitatively more pronounced in the APP^SAA^ genotype (Supplementary Fig. S1A, B). In addition, we also observe little difference in the time spent in the center of the OFT, suggesting that there is not an overt, age-dependent increase in anxiety-like behaviors (Supplementary Fig. S1C, D). The lack of evident locomotor impairment is also observed in an array of motor function testing. Each genotype displayed equivalent performance in sensorimotor function during nasal adhesive removal, strength in the wirehang test, and limb rigidity at 12 months of age (Supplementary Fig. S2A-C). Qualitatively in line with other neurologically intact mouse models^18^, this suggests that neither APP-KI genotype imparts greater dysfunction in these behaviors. At 12mo of age, APP^SAA^ mice displayed increased fecal output, but total intestinal transit was identical between the two strains tested (Supplementary Fig. S2D-E), suggesting that there may be some mild GI effects in the presence of amyloid pathology.

To characterize cognitive behaviors, we tested both genotypes longitudinally between 2 and 12mo of age in a small battery of cognitive tests. Using the object location test (OLT), neither genotype displayed a progressive loss of spatial memory in this test across age. While the APP^SAA^ genotype maintained a consistent discrimination of the object in a novel location (Fig. 1A), the APP^WT^ genotype consistently performed no better than chance (Fig. 1B). Similarly, in the Y-maze test of spatial working memory, we observe no progressive loss of spatial memory in either genotype through 12mo of age, with neither displaying any evidence of repetitive arm entries indicative of cognitive impairment (Fig. 1C, D). In the Barnes maze, both genotypes were able to quickly learn the location of the goal (likely during the initial habituation trial) at all ages tested (Fig. 1E, F). Following a 72hr retention interval, all animals were able to find the goal and consistently explore the goal quadrant, with no progressive loss of memory over time (Fig. 1G, J) While the mixed sex cohorts used in the present study were not sufficiently powered to detect sex differences, no overt differences between sexes were observed in any of the behavioral tests performed.

**Figure 1.**
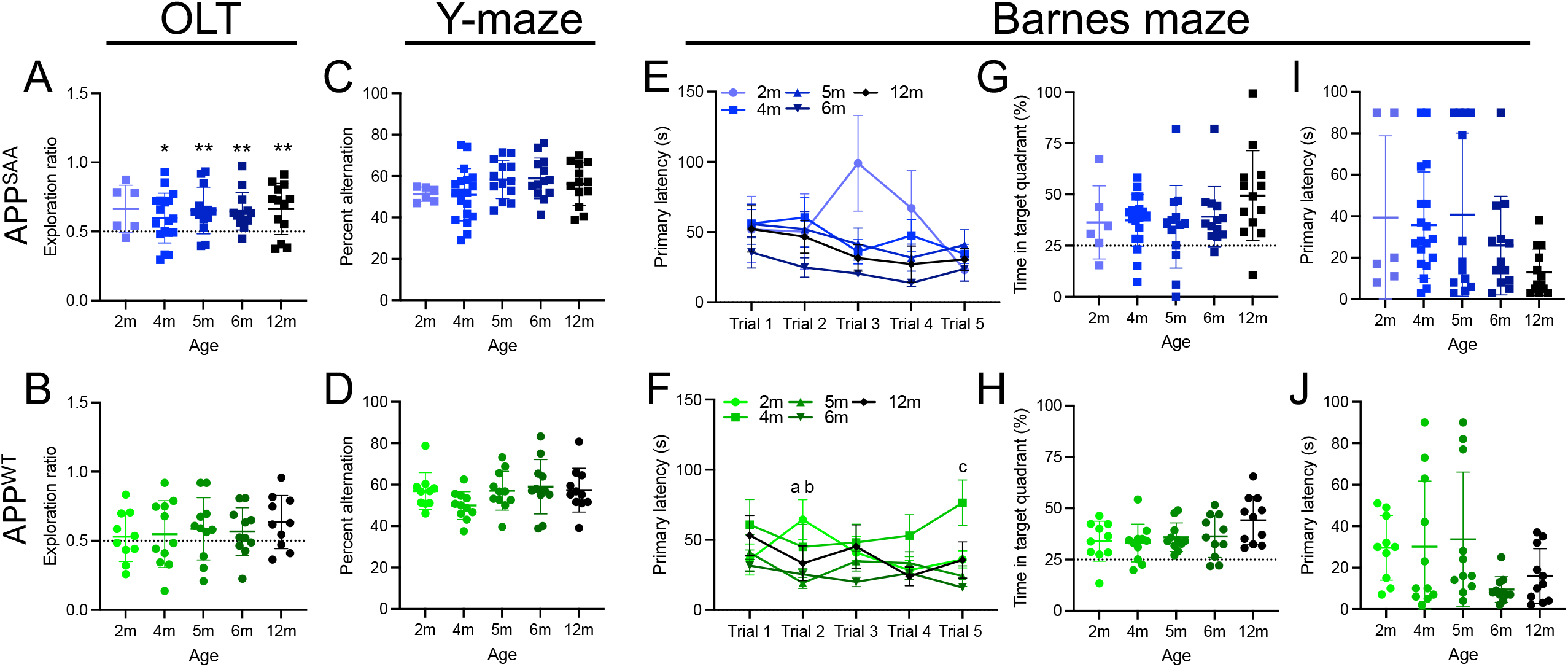
APP-KI mice do not show progressive cognitive impairment through 12 months of age. Male and female APP^SAA^ (**A, C, E, G, I**) and APP^WT^ (**B, D, F, H, J**) mice were tested from 2 through 12month (m) of age, at the indicated time points across a battery of cognitive behavior tests. **A, B)** Exploration ratio in the object location test (OLT). **C, D)** Percent alternation in the Y-maze. **E-J)** Barnes maze performance with longitudinal training period (**E, F**), followed by a 72hr probe trial measuring the time in target quadrant (**G, H**) and the primary latency to target (**I, J**). Points represent individuals (excluding **E, F**, where points represent the group mean), bars represent the mean and SEM. N= 10-11 APP^WT^ and 6-19 APP^SAA^. Data analyzed using mixed effects model with repeated measures and Dunnett’s multiple comparisons test to compare each age to 2m timepoint in **C-J**. \**p*≤0.05; \*\**p*≤0.01; \*\*\**p*≤0.001; \*\*\*\**p*≤0.0001; a,b,c *p*≤0.05 2m compared to (a) 5m, (b) 6m, (c) 4m.

**Figure 2.**
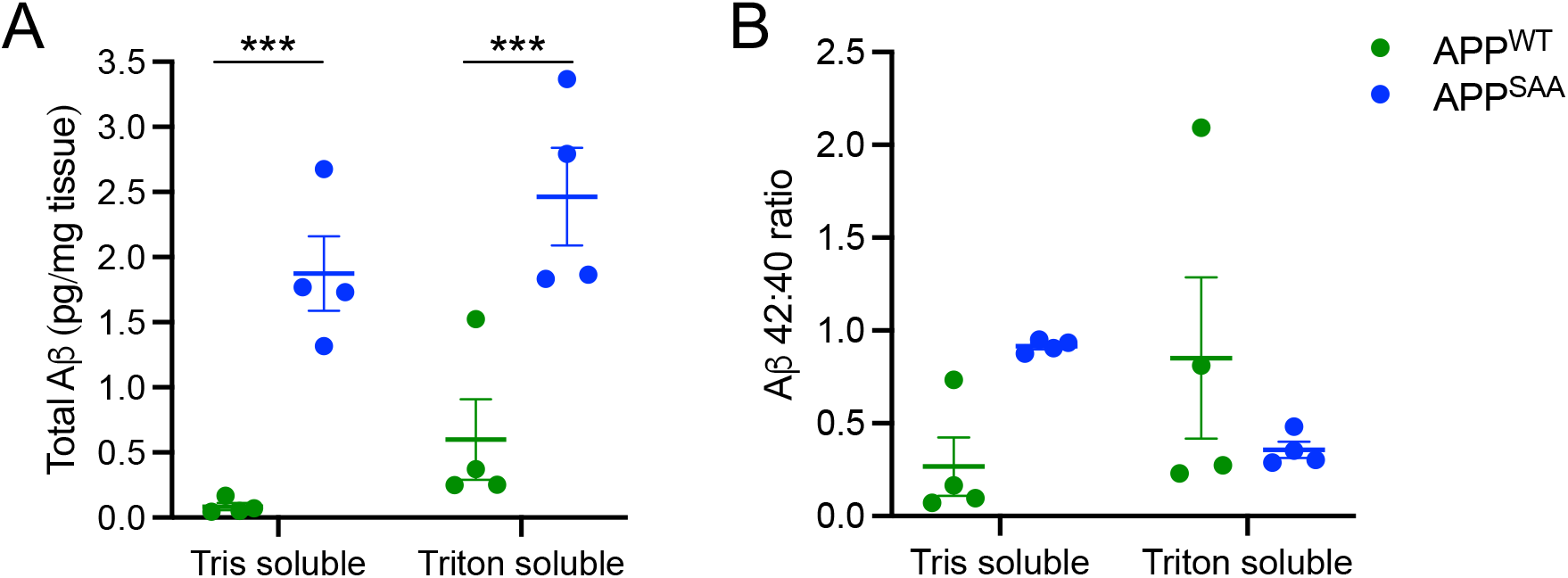
APP-KI mice produce human amyloid beta. Tris soluble and triton soluble hippocampal protein from 12-month-old male and female APP^SAA^ and APP^WT^ mice were analyzed by multiplex ELISA. **A)** Total Aβ including Aβ38, Aβ40, and Aβ42 **B)** Aβ 42:40 ratio. A significant genotype by solubility interaction effect was observed in Aβ 42:40 ratio (p=0.0297). Points represent individuals, bars represent the mean and SEM. N=4. Data analyzed by 2-way ANOVA with Tukey’s post-hoc tests comparing each genotype. \*\*\**p*≤0.001.

To confirm the presence of increased human amyloid beta (Aβ), we performed detergent fractionation of brain tissue lysates from each genotype at 12mo of age. In line with prior reports^15^, we observed the presence of both detergent soluble and insoluble human Aβ by ELISA, with generally increased levels in the APP^SAA^ genotype (Fig. 3A, B). Given the behavioral outcomes, we conclude that the presence of amyloid burden in these strains is not associated with cognitive impairment in these hallmark behaviors.

## Discussion

Knock-in models, such as the APP-KI strains used in this study, represent an important tool to understand etiological mechanisms that underlie amyloid pathologies and therapeutic interventions aimed at their clearance. By removing the presence of the murine amyloid ortholog, potential confounding interactions between the human and murine amyloids are avoided. Transcriptional control by the native human promoter in these models further allows a clearer understanding of how amyloid proteins respond to disease-relevant insults, such as immune modulation, metabolic input, or environmental exposures. Therefore, it is critical to evaluate baseline pathologies and behaviors in these model systems to have a foundation to explore such perturbations.

The APP-KI mice used in this study are one of these emerging and recently described model systems and currently lack complete characterization. While prior studies have characterized amyloid pathology and neuroinflammation in these two models^15^, to our knowledge, there is no published dataset of their cognitive behaviors. We therefore set out to identify a behavioral “tipping point,” an age where these genotypes began to show cognitive impairment or an age where the more pathogenic APP^SAA^ genotype separated behaviorally from the APP^WT^ genotype. Such a timepoint is important for timing potential interventions that seek to accelerate or diminish disease outcomes, directly test etiological contributions, or evaluate therapeutic interventions.

APP^SAA^ mice have been reported to display progressive amyloid pathology and neuroinflammation starting at 4mo of age^15^. We therefore hypothesized that these mice would display cognitive impairment as pathology progressed. However, despite assessing behaviors longitudinally for 12mo and confirming the presence of human Aβ accumulation in both mouse models, we were unable to detect progressive cognitive impairment in either genotype. Of note, the APP^WT^ mice preformed at chance levels on the OLT at all timepoints, suggesting a possible cognitive impairment even in young mice. However, lack of observable cognitive deficits on the other cognitive tests, suggests that this deficit is either very specific to the type of short-term object location memory tested, or is due to a confounding variable such as differences in motivation, hyperactivity, or visual acuity. Indeed, Xia et al. have reported age-related hyperactivity in APP^SAA^ mice that may similarly occur in the APP^WT^ genotype^15^. However, we did not observe any age-related increase in hyperactivity in the open field test for either genotype, possibly masked by repeat testing.

Interestingly, we did observe a genotype effect in aspects of gastrointestinal functions-whereby APP^SAA^ mice displayed increased fecal output in comparison to APP^WT^ mice. While there was no difference in total intestinal transit, this suggests that further evaluation of GI functions in this model system is warranted. GI disturbances are common in individuals living with PD and LBD, often present with Aβ co-pathologies. This may highlight the potential utility of these mouse models to explore the association between amyloid pathology and GI function in broader contexts.

We appreciate that murine behaviors have a number of caveats, including environmental variability, however the behavioral tests utilized in this study have been used to identify cognitive impairments in a range of AD models ^19^. For example, out of a battery of cognitive tests, the Barnes Maze test was found to be the most sensitive for detecting the development of cognitive impairment in the overexpression based 3xTg amyloid model ^20^. Further, while the longitudinal design of the present study provides the ability to observe disease progression, repeated testing may also result in habituation which may mask more subtle outcomes in these tests. Although no cognitive deficits were seen at 12 mo of age, it is possible that behavioral deficits may appear at later timepoints, as amyloid pathology continues to progress. Indeed, some amyloid-dependent models of AD do not display observable cognitive impairment until much later in age ^21,22^. It is also possible that the total amount of amyloid accumulation may not be sufficient to drive cognitive impairment. While many amyloid based models display cognitive impairment, increasing evidence suggests that Aβ pathology alone may not be sufficient to replicate certain aspects of human AD. Co-pathology with neurofibrillary tangles, synapse loss, and neuroinflammation may instead be more accurate predictors of cognitive performance ^23^.

No AD mouse model can fully recapitulate the human condition; however, when properly characterized, model systems are essential tools to address specific biological and translational questions. The present study provides a longitudinal characterization of the behavioral phenotypes of APP^SAA^ and APP^WT^ mice in a mixed-sex cohort over the course of a year, providing essential baseline behavioral data to inform future work. While we do not observe progressive cognitive impairment during this time period, additional perturbations and gene-by-environment interactions may be necessary to trigger cognitive decline in these models. Future studies into how various insults shape pathological and behavioral outcomes in these relevant Aβ-dependent mouse models will provide substantial insights into how various factors interact to promote disease.

## Acknowledgments

We thank David Weinshenker and members of the Sampson lab for productive discussions, and Srikant Rangaraju for the initial mouse lines used in this study. We acknowledge support from the Emory Division of Animal Resources (DAR) at Emory University - Integrated Core Facilities of the Emory University School of Medicine and supported by the Georgia Clinical and Translational Science Alliance of the NIH (UL1TR002378). Funding for this work was derived from NIH/NIA F31AG076332 (LBR) and NIH/NIEHS R01ES032440 (TRS).

## Supplementary Figure Legends

**Supplementary Figure S1.**
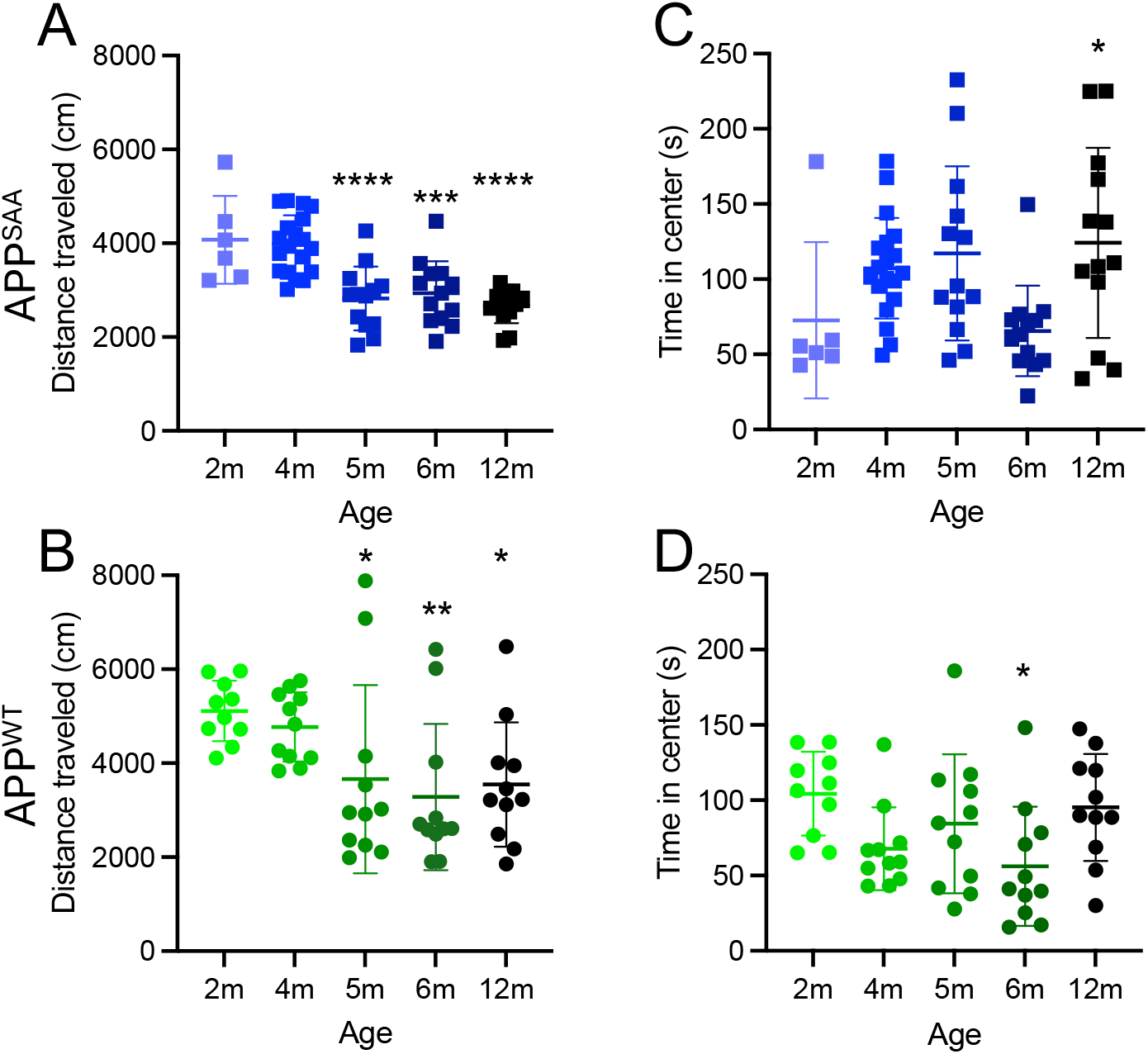
Performance of APP-KI mice in open field test through 12 months of age. Male and female APP^SAA^ (**A, C**) and APP^WT^ (**B, D**) mice were tested on the open field test for 10 minutes to assess motor and anxiety behaviors. Points represent individuals, bars represent the mean and SEM. N= 10-11 APP^WT^ and 6-19 APP^SAA^. Data analyzed using mixed effects model with repeated measures and Dunnett’s multiple comparisons test to compare each age to 2-month timepoint \**p*≤0.05; \*\**p*≤0.01; \*\*\**p*≤0.001; \*\*\*\**p*≤0.0001.

**Supplementary Fig. S2.**
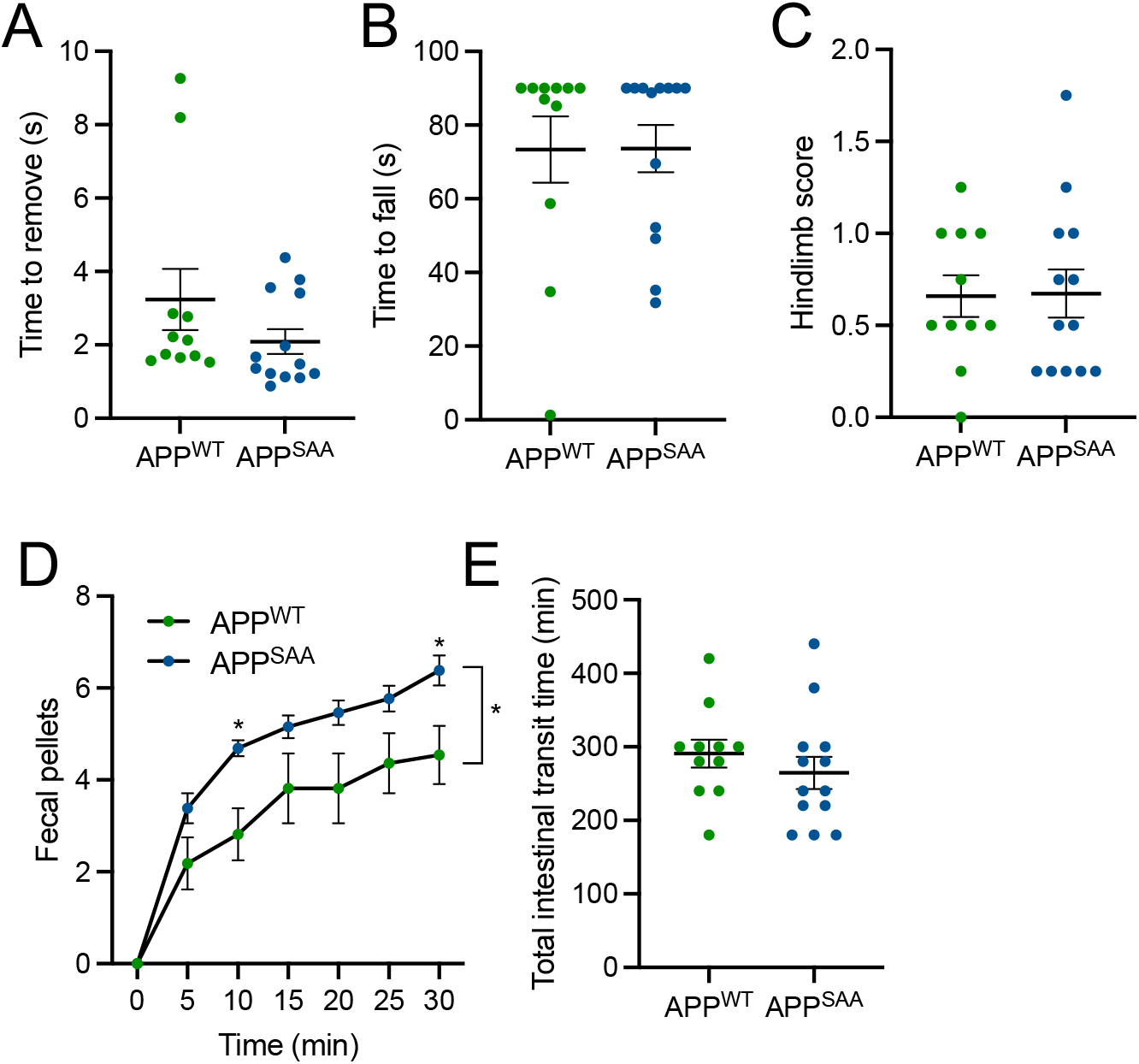
Performance of APP-KI mice in motor and gastrointestinal functions. Male and female APP^SAA^ and APP^WT^ mice were assayed at 12 months of age in tests of motor function.**A**) Time to remove a nasal adhesive. **B)** Time to fall on the wire hang test. **C)** Hindlimb rigidity score. **D)** Fecal output measured over 30 minutes. **E)** Total intestinal transit time as measured by carmine red elution. Points represent individuals (excluding **D** where points represent the group mean), bars represent the mean and SEM. N= 11-13. Data analyzed by unpaired T-tests in **A-C, E** and a repeated measures 2-way ANOVA with Sidek’s post-hoc test in **D**. \**p*≤0.05; \*\**p*≤0.01; \*\*\**p*≤0.001; \*\*\*\**p*≤0.0001

